# Fis connects two sensory pathways, quorum sensing and surface sensing, to control motility in *Vibrio parahaemolyticus*

**DOI:** 10.1101/2021.01.12.426476

**Authors:** Tague J.G., A. Regmi, G.J. Gregory, E.F. Boyd

## Abstract

Fis (<u>F</u>actor for <u>I</u>nversion <u>S</u>timulation) is a global regulator that is highly expressed during exponential growth and undetectable in stationary growth. Quorum sensing (QS) is a global regulatory mechanism that controls gene expression in response to cell density and growth phase. In *V. parahaemolyticus*, a marine species and a significant human pathogen, the QS regulatory sRNAs, Qrr1 to Qrr5, negatively regulate the high cell density QS master regulator OpaR. OpaR is a positive regulator of capsule polysaccharide (CPS) formation required for biofilm formation and a repressor of swarming motility. In *Vibrio parahaemolyticus*, we showed, using genetics and DNA binding assays, that Fis bound directly to the regulatory regions of the *qrr* genes and was a positive regulator of these genes. In the Δ*fis* mutant, *opaR* expression was induced and a robust CPS and biofilm was produced, while swarming motility was abolished. Expression analysis and promoter binding assays showed that Fis was a direct activator of both the lateral flagellum *laf* operon and the surface sensing *scrABC* operon, both required for swarming motility. In *in vitro* growth competition assays, Δ*fis* was outcompeted by wild type in minimal media supplemented with intestinal mucus, and we showed that Fis directly modulated catabolism gene expression. In *in vivo* colonization competition assays, Δ*fis* was outcompeted by wild type, indicating Fis is required for fitness. Overall, these data demonstrate a direct role for Fis in QS, motility, and metabolism in *V. parahaemolyticus*.

**IMPORTANCE:** In this study, we examined the role of Fis in modulating expression of the five-quorum sensing regulatory sRNAs, *qrr1* to *qrr5*, and showed that Fis is a direct positive regulator of QS, which oppositely controls CPS and swarming motility in *V. parahaemolyticus*. The Δ*fis* deletion mutant was swarming defective due to a requirement for Fis in lateral flagella and surface sensing gene expression. Thus, Fis links QS and surface sensing to control swarming motility and, indirectly, CPS production. Fis was also required for cell metabolism, acting as a direct regulator of several carbon catabolism loci. Both *in vitro* and *in vivo* competition assays showed that the Δ*fis* mutant had a significant defect compared to wild type. Overall, our data demonstrates that Fis plays a critical role in *V. parahaemolyticus* physiology that was previously unexamined.

## INTRODUCTION

The factor for inversion stimulation (Fis) is a nucleoid associated protein (NAP) that has two major functions in bacteria, chromosome organization and gene regulation (1, 2). Fis, along with other NAPs, is an important positive regulator of ribosome, tRNA and rRNA expression (3–6). As a transcriptional regulator, it can act as both an activator and repressor of a large number of genes (7–10). As an activator, Fis can directly bind to RNA polymerase (RNAP) to affect transcription, or indirectly control transcription via DNA supercoiling at promoters (3, 4, 11, 12). Fis controls DNA topology by regulating DNA gyrases (*gyrA* and *gyrB*) and DNA topoisomerase I (*topA*), required for DNA negative supercoiling in *Escherichia coli* and *Salmonella enterica* (7, 13, 14). In enteric species, Fis was shown to be a global regulator that responded to growth phases and abiotic stresses (8, 9, 15–22). In *E. coli*, Fis was highly expressed in early exponential phase cells and absent in stationary phase cells, under aerobic growth conditions (23, 24).

In *Vibrio cholerae,* it has been shown that Fis controls the quorum sensing (QS) regulatory sRNAs (25). In this species, it was proposed that Fis acted at the level of LuxO, the QS response regulator and a sigma factor-54 dependent activator, both required for transcription of the regulatory sRNAs *qrr1* to *qrr4*. In *V. cholerae*, the Qrrs repress the QS high cell density (HCD) regulator HapR and activate the QS low cell density regulator AphA. In a Δ*fis* mutant in this species, the *qrr* genes were repressed and *hapR* was expressed at wild type levels (25). In *E. coli* and *Salmonella enterica*, Fis was shown to control virulence, motility, and metabolism (9, 10, 16, 26). These studies identified 100s of genes whose expression *in vivo* is either enhanced or repressed by Fis. In *S. enterica,* the polar flagellum genes and genes within several pathogenicity islands were differentially expressed between a Δ*fis* mutant and wild type. In *Dickeya zeae*, a plant pathogen, swimming and swarming motility was reduced in a Δ*fis* mutant strain and a total of 490 genes were significantly regulated by Fis, including genes involved in the QS pathway, metabolic pathways, and capsule polysaccharide (CPS) production, amongst others (21, 27–29).

*Vibrio parahaemolyticus* is the leading cause of bacterial seafood-borne gastroenteritis worldwide and in the United States alone, tens of thousands of cases of *V. parahaemolyticus* infections are reported each year (30–33). A *V. parahaemolyticus* infection causes inflammatory diarrhea and its main virulence factors are two type three secretions systems and their effector proteins (34–36). Unlike *V. cholerae* that only produces a single polar flagellum, *V. parahaemolyticus* produces both a polar flagellum and lateral flagella expressed from the Flh (Fli) and Laf loci, respectively (37–39). The polar flagellum, required for swimming motility, is produced in cells grown in liquid media and is under the control of sigma factors RpoN (σ54) and FliAP (σ28). The lateral flagella, required for swarming motility, are produced on solid media and are under the control of RpoN, LafK a σ54-dependent regulator, and a second σ28 factor, FliAL (38, 40). Disruption of σ54 abolishes all motility, whereas deletion of the two σ28 sigma factors, FliAP (FliA) and FliAL, abolishes swimming and swarming, respectively (38, 41). Overall, control of motility in the dual flagellar system of *V. parahaemolyticus* differs significantly from monoflagellar systems of enteric species (42–44).

Previously, it was demonstrated that bacterial motility and metabolism require a functional QS pathway in *V. parahaemolyticus* (45). It was shown that a Δ*luxO* deletion mutant, in which the *qrr* sRNAs are repressed, constitutively expressed *opaR* (the *hapR* homolog). The Δ*luxO* mutant had reduced swimming motility, but swarming motility was abolished, while a Δ*opaR* deletion mutant was hyper-motile and swarming proficient (45). Additionally, studies have shown that the *V. parahaemolyticus scrABC* surface sensing operon activates swarming motility and represses capsular polysaccharide (CPS) formation by reducing the cellular levels of c-di-GMP (46–48). A deletion of the *scrABC* operon induces high c-di-GMP levels that repress *laf* and induce *cps* gene expression (46–48). The QS regulator, OpaR, is a direct repressor of *laf*, required for swarming, and an activator of the *cps* operon, required for CPS formation required for biofilm formation (49).

Here, we characterized the role of Fis in the gastrointestinal pathogen *V. parahaemolyticus*, and show that Fis connects the QS and surface sensing signaling pathways in this species. We determined the expression pattern of *fis* across the growth curve and constructed an in-frame Δ*fis* deletion mutant to examine its role in *V.* *parahaemolyticus* physiology. We examined the role of Fis in the QS pathway, specifically its control of the five regulatory sRNAs, *qrr1* to *qrr5*, using transcriptional GFP reporter assays and DNA binding analyses. The effects of a *fis* deletion on swimming and swarming motility was determined and the mechanism for the requirement for Fis in swarming motility was uncovered. To investigate whether Fis plays additional roles in *V. parahaemolyticus* physiology, we performed *in vitro* growth competition assays between the Δ*fis* mutant and a *lacZ* knock-in WT strain, WBWlacZ. Further, GFP reporters and DNA binding assays were performed to examine a direct role for Fis in carbon metabolism. *In vivo* colonization competition assays were also performed using a streptomycin-pretreated adult mouse model of colonization. This study demonstrates that Fis integrates the QS and surface sensing pathways to control swarming motility and is important for overall cell function.

## RESULTS

### *fis* expression is controlled in a growth dependent manner

Locus tag VP2885 is annotated as a Fis protein homolog, a 98 amino acid protein that shows 100% protein identity with Fis from *V. cholerae* and 82% protein identity with Fis from *E. coli*. Fis is an abundant protein in *E. coli*, highly expressed in exponential phase cells. In *V. parahaemolyticus* RIMD2210633, we determined the expression pattern of *fis* across the growth curve, via RNA isolated from wild type cells grown in LB 3% NaCl (LBS) at 37°C aerobically at various optical densities (ODs). Using quantitative real time PCR (qPCR) analysis, *fis* showed highest expression levels in exponential cells at ODs 0.15, 0.25, and 0.5 and then rapidly declined at ODs 0.8 and 1.0, as cells entered stationary phase (**Fig. S1**). These data show that Fis in *V. parahaemolyticus* has a similar expression pattern to Fis in *E. coli* and also what has been demonstrated in *V. cholerae* (2, 23, 25).

### Fis positively regulates *qrr* sRNAs

To determine the role of Fis in the regulation of *qrr1* to *qrr5* in *V. parahaemolyticus*, we identified putative Fis binding sites within the regulatory regions of all five sRNAs. To confirm Fis binding, we purified the Fis protein and constructed DNA probes of the regulatory region of each *qrr* to perform electrophoretic mobility shift assays (EMSA) with increasing concentrations of purified Fis. Direct binding was shown in a concentration dependent manner in all five P*qrr-*Fis EMSAs (**Fig. 1A-E**). A fragment of DNA with no putative Fis binding site was used as a non-binding control **(Fig. S2A)** and the regulatory region of *gyrA* was used as a Fis binding positive control (**Fig. S2B**). The regulatory region of *qrr1* to *qrr5* contains between one and three putative binding sites, which may explain the differences in intensity observed in the EMSAs (**Fig. 2**). P*qrr3* only contained one putative Fis binding site and showed fewer shifts in the gel compared to P*qrr2,* which contains three putative Fis binding sites.

**FIG 1.**
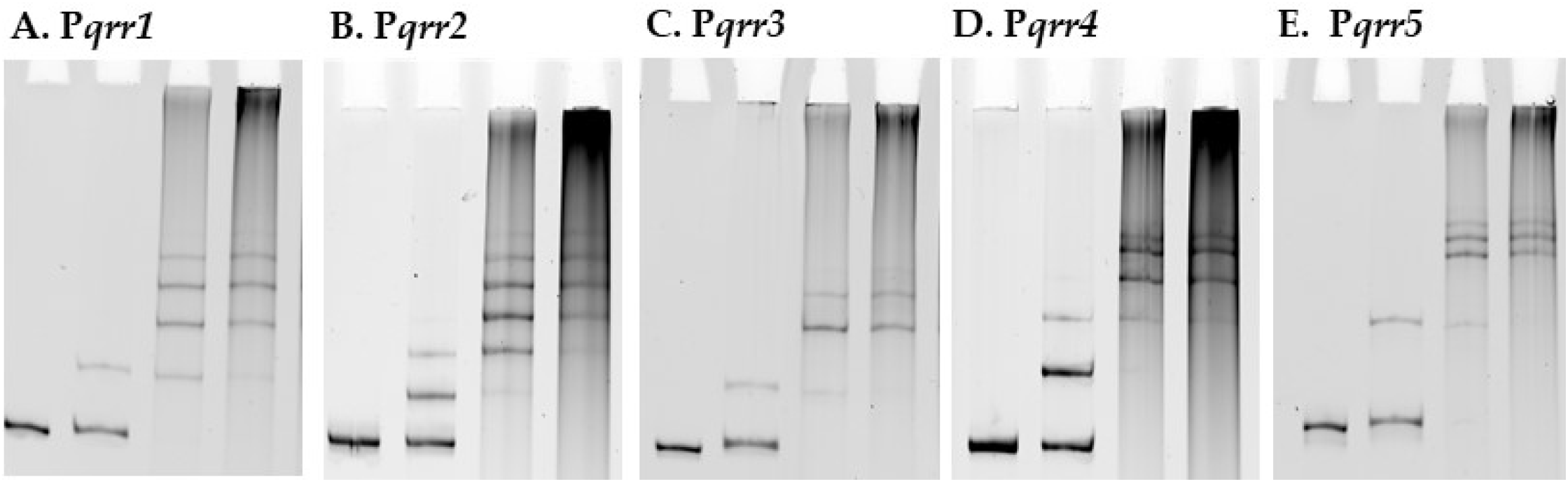
Fis binds to the regulatory regions of *V. parahaemolyticus* regulatory sRNAs *qrr1* to *qrr5.* **A-E.** Electrophoretic mobility shift assays using purified Fis and the regulatory regions of *qrr1* to *qrr5*. DNA:protein ratios are as follows: 1:0, 1:1, 1:20, 1:50.

**FIG 2.**
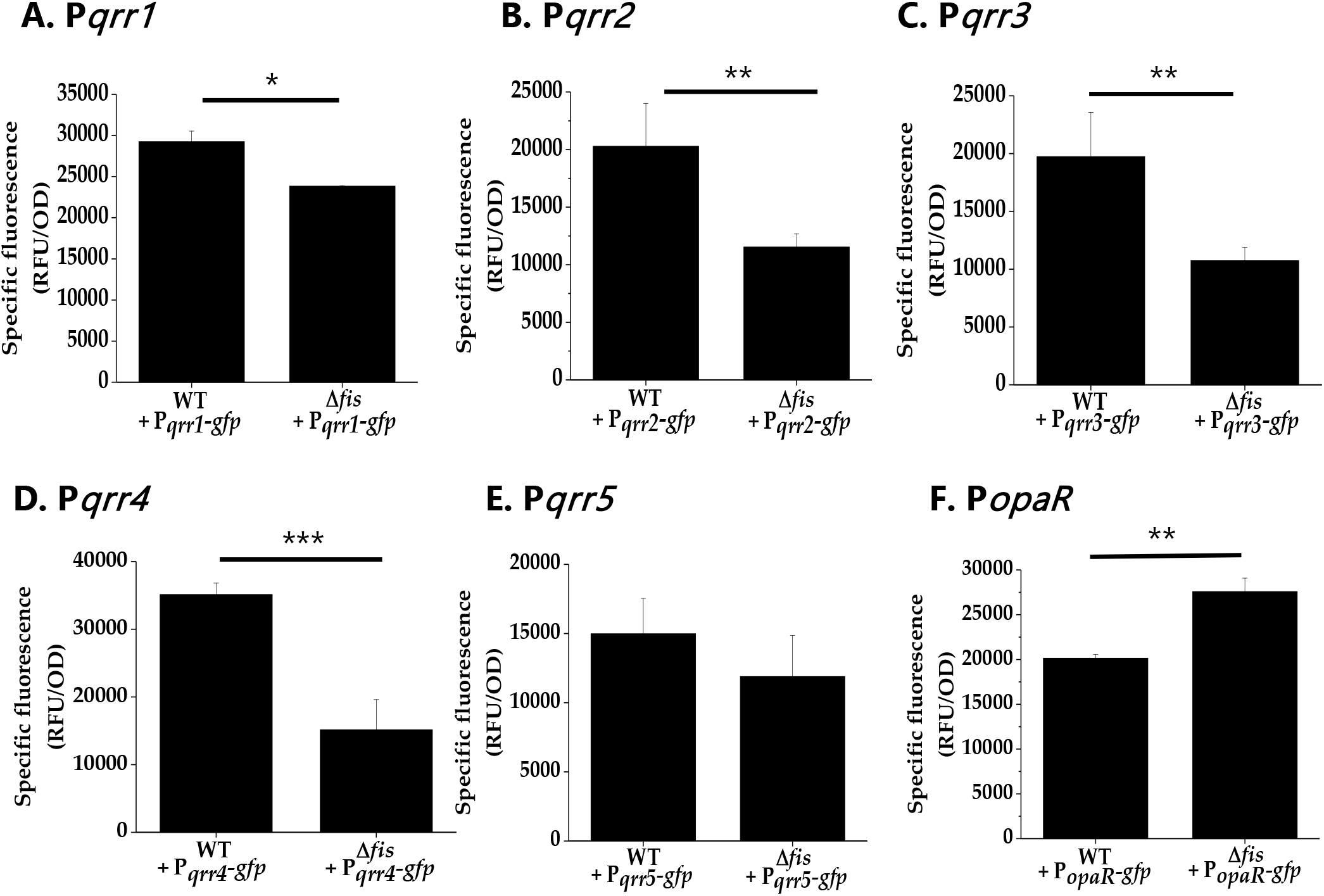
Fis is a positive regulator of the *qrr* genes. **A-F.** *qrr1* to *qrr5* GFP transcriptional reporter assays in wild type and the Δ*fis* mutant in cultures grown to OD 0.4-0.45 and measured for specific fluorescence (RFU/OD). Means and standard deviations of three biological replicates are plotted. Statistics calculated using a Student’s t-test. (***, P < 0.001).

To further characterize the role of Fis in *V. parahaemolyticus*, an in-frame deletion of *fis* was constructed by deleting 279-bp of VP2885. We examined growth of the Δ*fis* mutant in LBS broth and found that it grew identical to wild type (**Fig. S3A**). However, on LBS agar plates, the Δ*fis* mutant formed a small colony morphology compared to wild type, which was complemented with a functional copy of *fis* (**Fig. S3B & S3C**). To investigate whether Fis is a direct regulator of *qrr1* to *qrr5,* we performed green fluorescent protein (GFP) transcriptional reporter assays using the regulatory region of each *qrr*. Cells were grown to 0.4-0.45 OD and GFP levels measured using relative fluorescence normalized to OD (specific fluorescence). In Δ*fis*, the overall expression of P*qrr1*-*gfp* to P*qrr4*-*gfp* was significantly downregulated compared to wild type, indicating that Fis is a direct positive regulator of *qrr1* to *qrr4* in *V. parahaemolyticus* (**Fig 2A-D**). Under the conditions examined, we observed a reduction in P*qrr5*-*gfp* expression in the Δ*fis* mutant relative to wild type but this reduction was not statistically significant (**Fig. 2E**).

Next, we examined whether *opaR* expression was changed in the Δ*fis* mutant using GFP reporter expression assays under the control of the *opaR* regulatory region, P*opaR*-*gfp*. Induction of P*opaR*-*gfp* activity was observed in the Δ*fis* mutant compared to wild type, indicating derepression of *opaR* (**Fig. 2F**). Examination of CPS, which manifests as a rough wrinkly colony morphology and is positively regulated by OpaR, demonstrated that the Δ*fis* mutant produced robust CPS and biofilm phenotypes, further supporting that *opaR* expression is induced in a Δ*fis* mutant (**Fig. S4**). In contrast, the Δ*opaR* mutant formed a smooth colony morphology indicating CPS is lacking (**Fig. S4A**).

### Fis is essential for motility in *V. parahaemolyticus*

We examined whether deletion of *fis* affected motility in *V. parahaemolyticus,* a species that produces both polar and lateral flagella. Swimming assays demonstrated that the Δ*fis* mutant had a defect in motility compared to wild type (**Fig. 3A**), while in swarming assays, motility was abolished (**Fig. 3B**). These data suggest that Fis is a positive regulator of motility, with an essential role in swarming motility. To confirm that both observed swimming and swarming defects are a result of the *fis* deletion, we complemented the Δ*fis* mutant with a functional copy of the *fis* gene under the control of an IPTG-inducible promoter. We observed rescue of both the swimming and swarming phenotypes in *fis* complemented strains (**Fig. S5A and S5B**).

**FIG 3.**
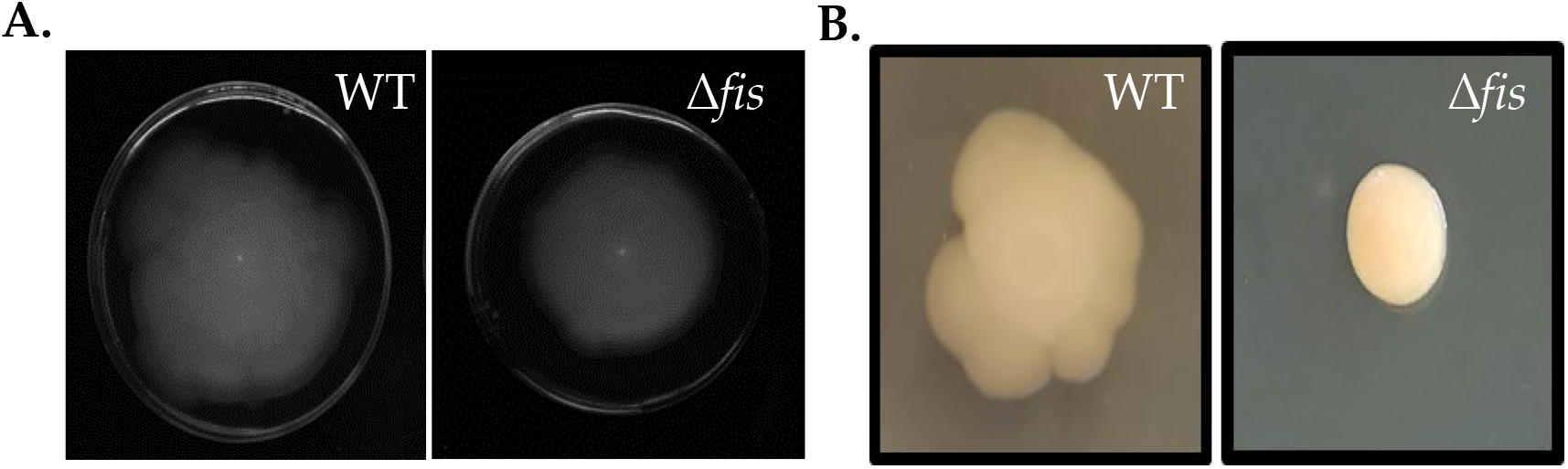
Motility defects in Δ*fis*. **A.** Swimming assays and **B.** swarming assays of *V. parahaemolyticus* wild type and the Δ*fis* mutant. All images are examples from three biological replicate.

In order to determine how Fis regulates swimming and swarming behavior, we used bioinformatics analysis and identified multiple putative Fis binding sites within the regulatory regions of both the polar and lateral flagella biosynthesis operons **(Fig. 4A and 4D)**. We performed EMSAs using purified Fis protein and DNA probes amplified from the regulatory regions of the polar flagellum operon (*flh* loci VP2235-PV2231) and the lateral flagellum operon (*laf* loci VPA1550-VPA1557) (**Fig. 4B and 4E**). EMSAs demonstrated that Fis binds to the DNA probes of both P*flhA* and P*lafB*. (**Fig. 4B and 4E**). GFP reporter expression assays of cells grown in LBS broth did not show differential expression of P_*flhA*-*gfp*_ in the Δ*fis* mutant relative to wild type (**Fig. 4C**). In a reporter assay of cells grown on heart-infusion plates, P_*lafB*-*gfp*_ was repressed in the Δ*fis* mutant relative to wild type (**Fig. 4F**). Overall, these data show that Fis directly modulates lateral flagella biosynthesis, and is a positive regulator of swarming motility

**FIG 4.**
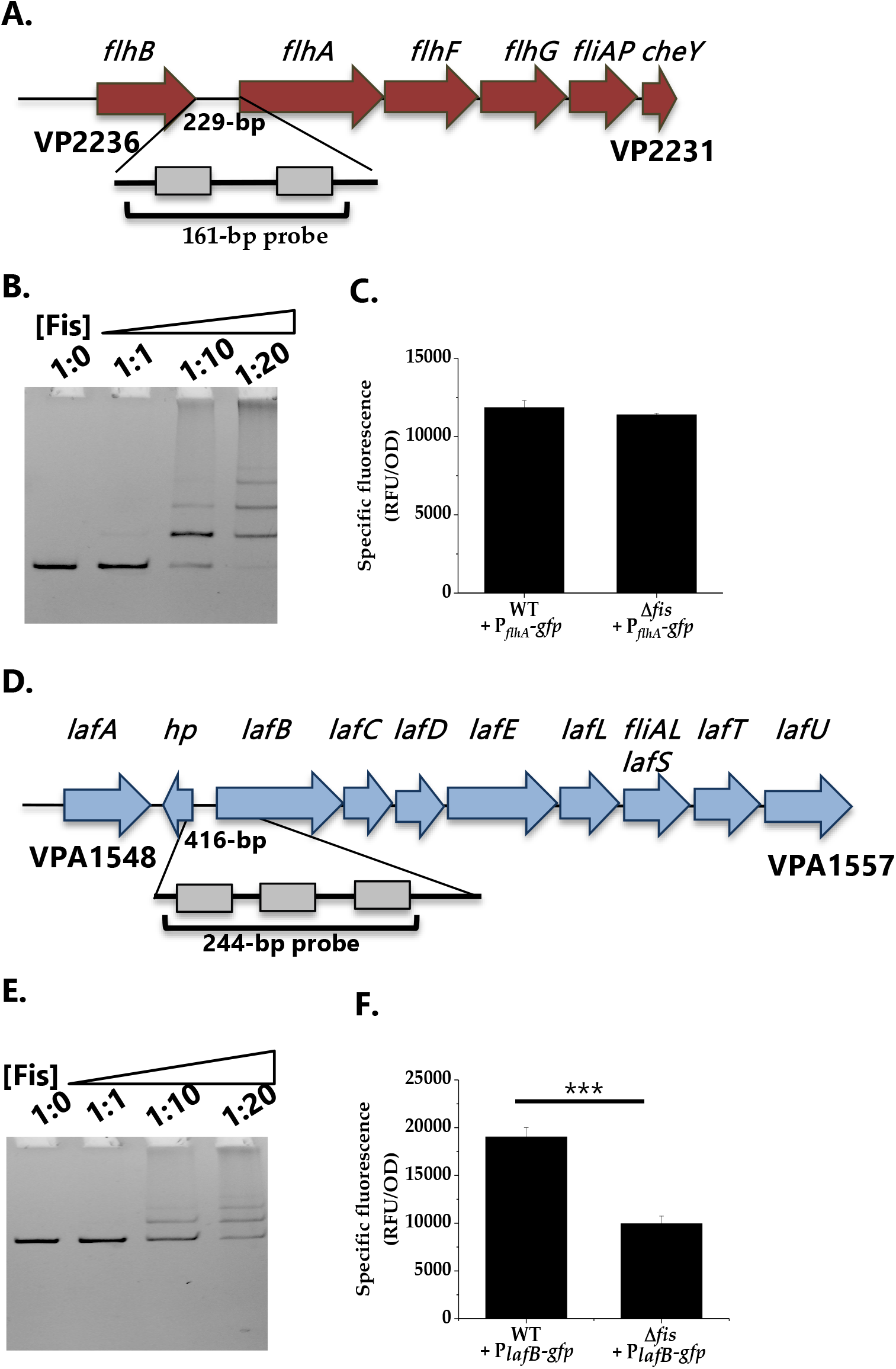
Regulation of polar and lateral flagella biosynthesis by Fis. **A.** Regulatory region of polar flagellum *flh* genes with putative Fis binding sites (BS) depicted as gray boxes. **B.** EMSA of Fis bound to P*flhA* in a concentration dependent manner. **C.** Transcriptional GFP reporter assay of P_*flhA*_-*gfp* in the Δ*fis* mutant relative to wild type. **D.** Regulatory region of lateral flagellum *laf* genes with putative Fis binding sites **E.** EMSA of Fis bound to P*laf* DNA probe. **F.** Transcriptional GFP reporter assay of P_*lafB*_-*gfp* between wild type and Δ*fis.* ***, P < 0.001.

### Fis is a positive regulator of the *scrABC* surface sensing operon

The *scrABC* operon has been shown to oppositely control swarming motility and CPS production, so we reasoned that Fis might also control this operon to co-ordinate with QS control of these phenotypes. First, we identified putative Fis binding sites in the regulatory region of *scrABC* (**Fig. 5A**), and then performed an EMSA, which demonstrated Fis binding to P*scrABC* in a concentration dependent manner (**Fig. 5B**). In GFP reporter assay, specific fluorescence was measured in both wild type and the Δ*fis* mutant harboring P_*scrABC*-*gfp*_ after growth to stationary phase on LB plates. This analysis showed that P_*scrABC*-*gfp*_ activity was significantly downregulated in the Δ*fis* mutant compared to wild type, indicating that Fis is a direct positive regulator of this operon (**Fig. 5C**). Overall, the data suggest that loss of swarming motility is due to Fis regulation of *scrABC* and the lateral flagella biosynthesis *laf* operon in *V. parahaemolyticus*.

**FIG 5.**
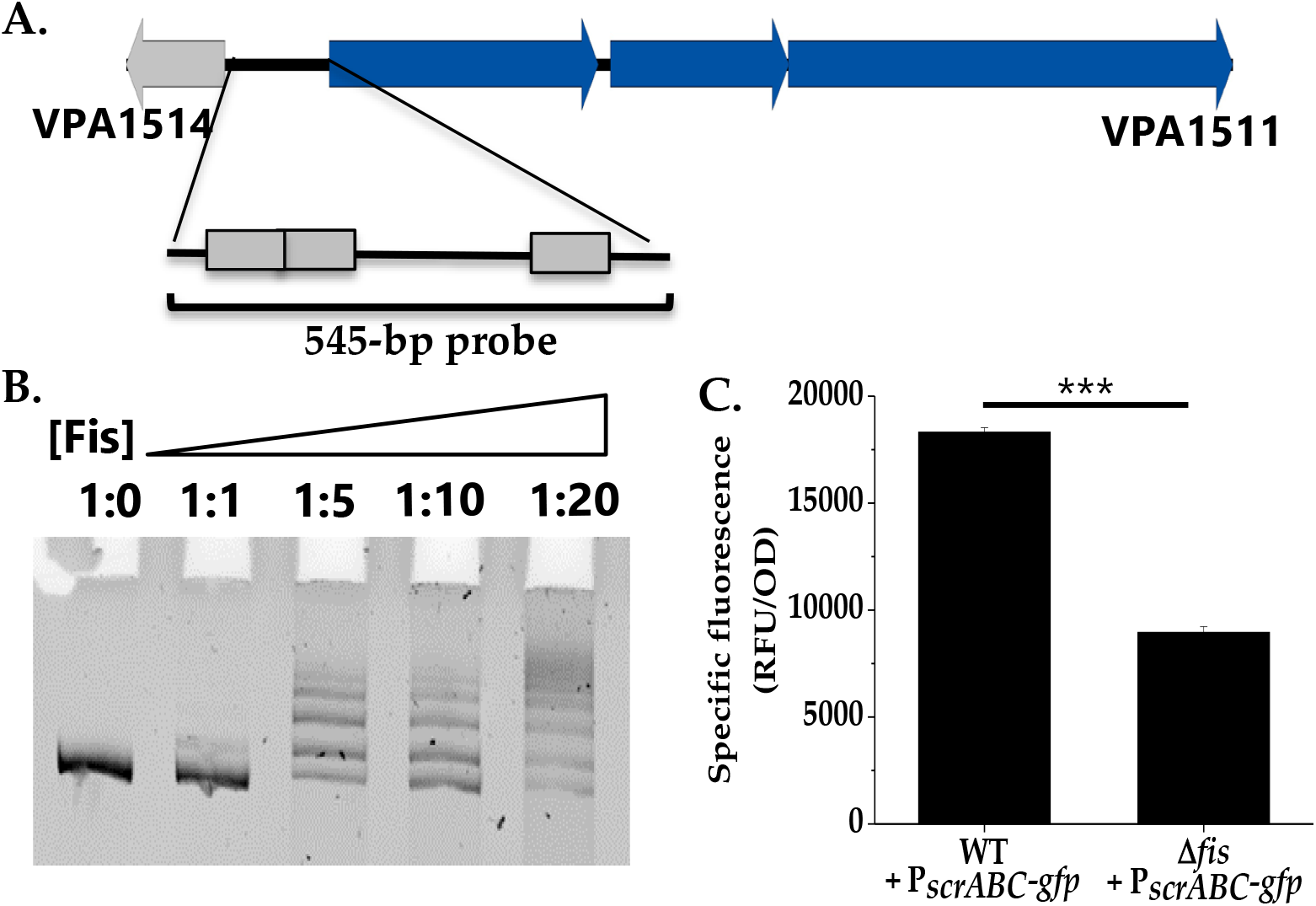
Fis is a positive regulator of the *scrABC* surface sensing operon. **A.** Putative Fis binding sites identified in the regulatory region of the *scrABC* surface sensing operon. **B.** An EMSA using purified Fis protein and the regulatory region of *scrABC* as a probe. **C.** GFP reporter assay of P_*scrABC*_-*gfp* in wild type and the Δ*fis* mutant. Specific fluorescence was calculated (RFU/OD) for three biological replicates and plotted as mean and standard deviation. Statistics were calculated using a Student’s t test (***, P < 0.001).

### Fis is a positive regulator of carbon catabolism gene clusters

To further investigate the role of Fis in *V. parahaemolyticus* physiology, we conducted growth competition assays between wild type and Δ*fis* in various carbon sources. For the *in vitro* competition assays, we used a β-galactosidase knock-in strain of RIMD2210633, strain WBWlacZ, which was previously demonstrated to behave identically to wild type in *in vitro* and *in vivo* studies (41, 45, 50, 51). *In vitro* growth competition assays were performed in M9 minimal media 3% NaCl (M9S) supplemented with mouse intestinal mucus as a sole carbon source. We also examined growth in individual carbon components of intestinal mucus. In the *in vitro* competition assays, Δ*fis* was significantly outcompeted by WBWlacZ in mouse intestinal mucus (CI 0.61), and mucus components L-arabinose (CI 0.4), D-glucosamine (CI 0.68) or D-gluconate (CI 0.78) (**Fig. 6**). These data suggest that Fis is an important regulator of cellular metabolism.

**FIG 6.**
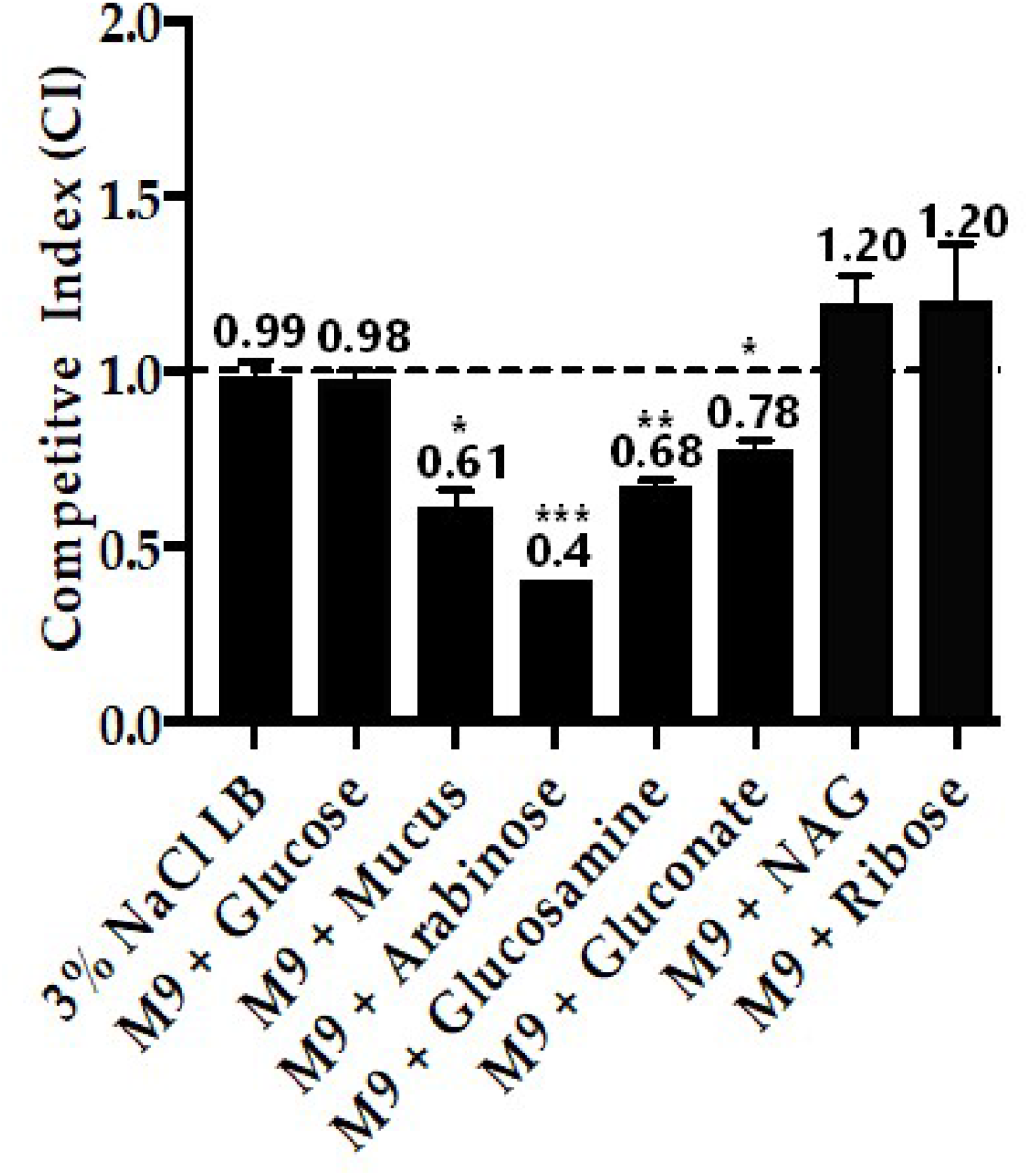
A. *In vitro* growth competition assay. WBWlacZ and Δ*fis* were grown in co-culture (1:1 ratio) for 24 hours in LBS, M9 supplemented with 100 μg/ml intestinal mucus, 10 mMof D-glucose, L-arabinose, D-glucosamine, D-gluconate, N-acetyl-D-glucosamine (NAG), or D-ribose. The assay was conducted in two biological replicates in triplicates. Error bar indicates SEM. Unpaired Student’s t test was conducted. The significant difference is denoted by asterisks (*, P<0.05, **, P<0.01,***, P<0.001).

Putative Fis binding sites were identified in the regulatory regions of *araBDAC* (L-arabinose catabolism) (**Fig. 7A**). EMSAs were performed using 138-bp and 152-bp DNA probes containing one and two putative Fis binding sites, respectively (**Fig. 7B**). In these assays, Fis bound to the regulatory region of *araBDAC*, and the binding was concentration dependent. In GFP transcriptional reporter assays, P_*araB*-*gfp*_ showed significantly lower activity in the Δ*fis* mutant compared to wild type, demonstrating that Fis is a direct activator of the *araBDAC* gene cluster. (**Fig. 7C**). Fis binding sites were also identified in the regulatory region of *gntK* (D-gluconate catabolism) (**Fig. 8A**). A DNA probe encompassing the *gntK* promoter region showed strong binding to Fis in an EMSA (**Fig. 8B**) and a GFP reporter assay of the regulatory region of *gntK* showed significantly lower expression levels in the Δ*fis* mutant compared to wild type (**Fig. 8C**). Fis binding sites were also identified in the regulatory region of *nagB* (D-glucosamine catabolism) (**Fig. 9A**). Fis bound to the regulatory region of *nagB* and showed significantly lower activity in the GFP reporter assay (**Fig. 9B and 9C)**. Overall, our results demonstrated that Fis is a direct positive regulator of the catabolism genes, *araB, gntK,* and *nagB*, in *V. parahaemolyticus*.

**FIG 7.**
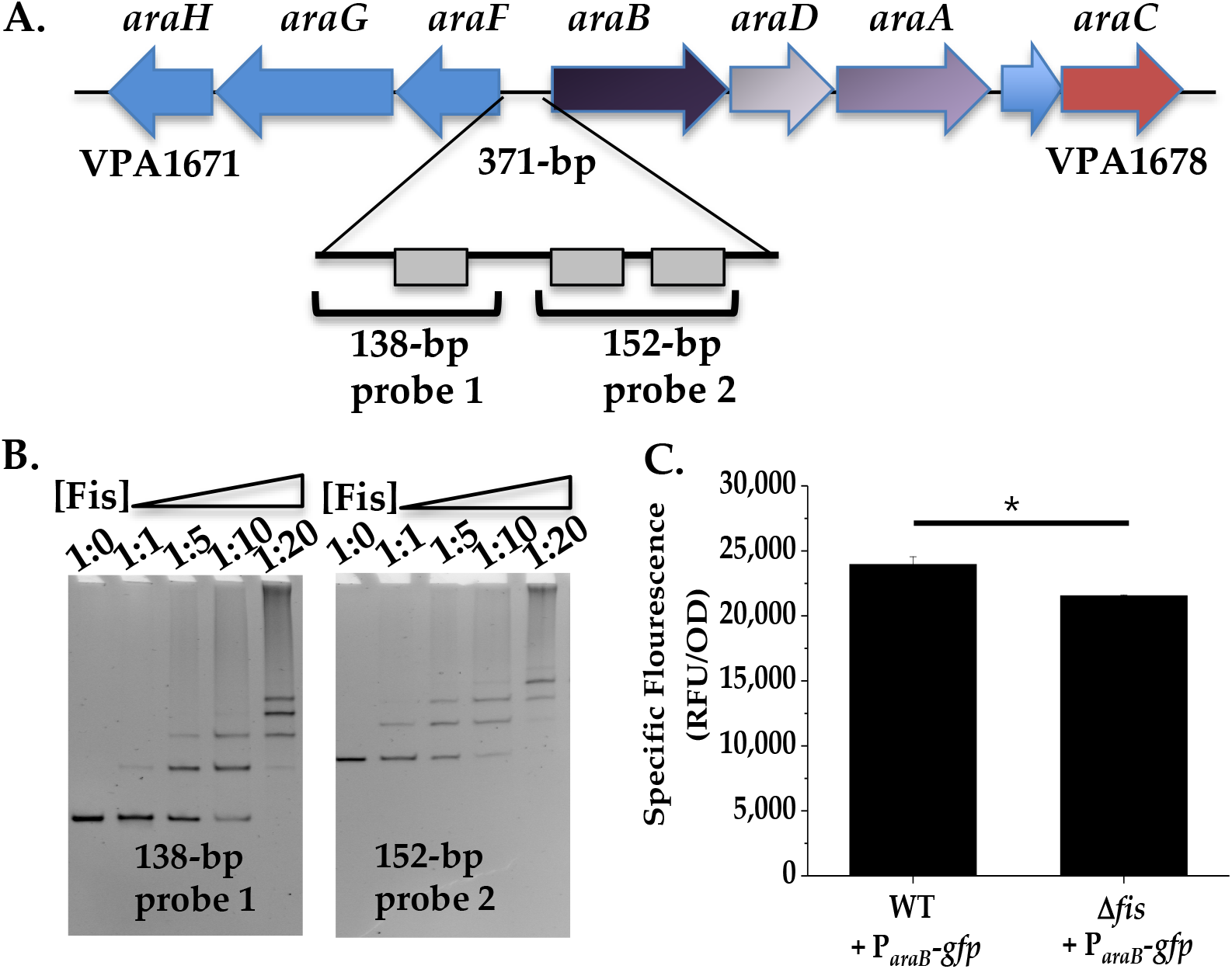
Regulation of *araBDAC* operon by Fis. **A.** Fis binding sites identified in the regulatory region of the *araBDAC* operon depicted as grey boxes. **B.** P*araB* was divided into two probes, 138-bp and 152-bp, each containing at least one putative Fis binding site. DNA probes were used in EMSAs with purified increasing ratios of DNA:Fis, and binding was observed as shift in the gel. **C.** GFP transcriptional reporter assay of the *araBDAC* regulatory region in wild type and the Δ*fis* mutant. Means and standard deviations of two biological replicates are shown. Statistics calculated using a Student’s t-test (*, P < 0.05).

**FIG 8.**
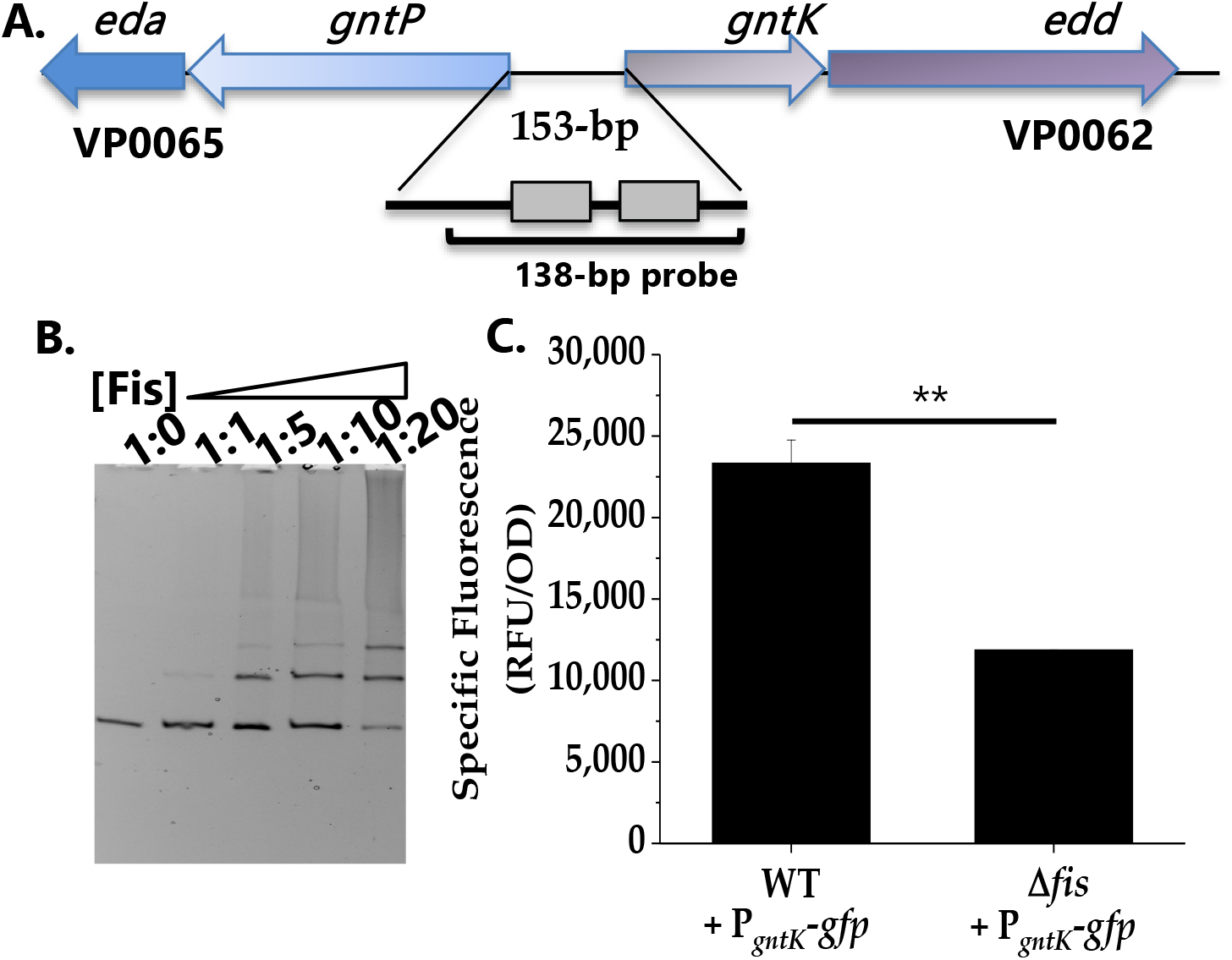
Regulation of *gntK* by Fis. **A.** Two putative Fis binding sites in the regulatory region of *gntK*. **B.** EMSA with purified Fis protein. **C.** GFP transcriptional reporter assay of *gntK* in wild type and Δ*fis*. Means and standard deviations of two biological replicates are shown. Statistics calculated using a Student’s t-test (**, P < 0.01).

**FIG 9.**
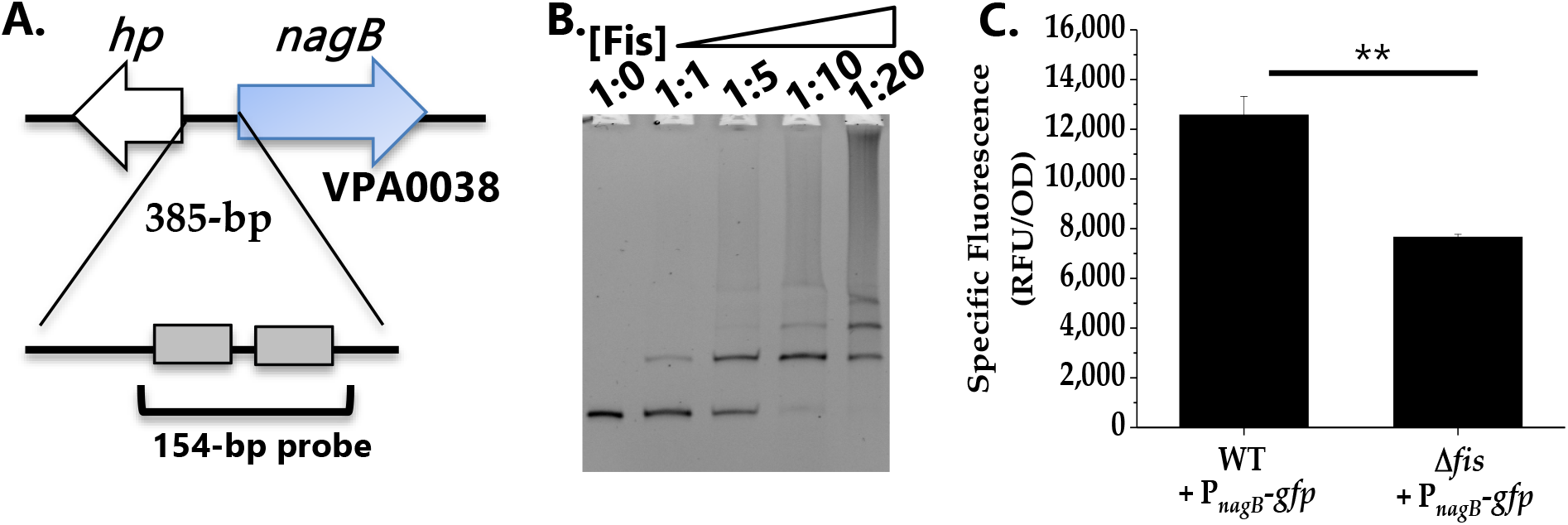
Regulation of *nagB* by Fis. **A.** Fis binding sites identified in the regulatory region of *nagB*. **B.** EMSA analysis of Fis binding to regulatory region of *nagB*. **C.** GFP transcriptional reporter assay with P_*nagB*_-*gfp*. Means and standard deviations of two biological replicates are shown. Statistics calculated using a Student’s t-test (**, P < 0.01).

### Fis is required for *in vivo* fitness in *V. parahaemolyticus*

To determine whether Fis contributes to *in vivo* fitness of *V. parahaemolyticus*, *in vivo* colonization competition assays were performed using a streptomycin pretreated adult mouse model of colonization (41, 50–52). WBWlacZ, is a *lacZ* positive strain that was previously demonstrated to behave similarly to wild type in *in vivo* and *in vitro* competition assays (41, 50–52). The *in vivo* competition assay determined the ability of WBWlacZ and the Δ*fis* mutant to co-colonize the intestinal tract of streptomycin pretreated adult mice. Competition assays were performed in adult C57BL/6 mice pretreated with an orogastric dose of streptomycin (20 mg/animal) 24 hrs prior to orogastric co-inoculation with a 1:1 mixture of *V. parahaemolyticus* WBWlacZ and Δ*fis* (n=9). In an *in vitro* assay in LB using the same inoculum, the WBWlacZ vs Δ*fis* had a CI of 1.17 whereas *in vivo* the WBWlacZ vs Δ*fis* the CI was 0.57 indicating that the mutant was significantly (*p* < 0.01) outcompeted by WBWlacZ (**Fig. S6**). This indicates that Δ*fis* has decreased fitness *in vivo* compared to the wild type strain.

## DISCUSSION

NAPs present in bacteria, such as Fis, bind and bend DNA to aid in DNA compaction, and are also important global regulators of gene expression. In *V. parahaemolyticus* we show, similar to enteric species, that *fis* is expressed in early to mid-exponential phase cells and declines in late exponential and stationary phase cells. Our work showed that Fis was a direct positive regulator of the QS regulatory sRNAs *qrr1* to *qrr5*, binding to the promoter region of each. Further, we showed that expression of the QS master regulator, *opaR,* was derepressed in the Δ*fis* mutant and the Δ*fis* mutant produced a robust CPS. Only one other study has shown a direct role of Fis in a QS pathway, which also showed that Fis activated the *qrr* sRNAs in *V. cholerae*. In this study, a Δ *fis* mutant showed HapR (OpaR homolog) constitutively expressed (25). In both *V. cholerae* and *V. parahaemolyticus*, *qrr* expression requires RNAP containing sigma-54 and the sigma-54 dependent activator LuxO to initiate transcription (45, 53). NAPs are known to play important roles in modulating sigma factor binding to promoter regions that aid or inhibit RNAP initiation of transcription (54). For example, in *E. coli*, expression of the major stationary phase binding protein Dps is repressed by Fis, ensuring repression in exponential phase but expression of the *dps* locus in stationary phase cells (55). Fis reduced transcription initiation by sigma-70-RNAP at the *dps* promoter but had no effect on sigma-38-RNAP, the stationary phase sigma factor (55). As stated above, *qrr* transcription requires sigma-54-RNAP, but unlike all other sigma factors, sigma-54 requires an additional factor, known as an enhancer binding protein (EBP), to activate transcription. EBPs bind upstream of the promoter and therefore require DNA binding and bending protein such as Fis, IHF, or H-NS to allow contact between the EBP and the RNAP holoenzyme (54). We speculate that the NAP Fis may play a key role in *qrr* expression by aiding sigma-54-RNAP interaction with EBP LuxO.

Studies have shown that although Fis may bind to a large number of regulatory regions, only a portion of these sites are significantly regulated by Fis (56, 57). For example, ChIP-seq analysis in *E. coli* showed 1464 Fis binding sites, but only 462 genes were differential regulated by Fis under the conditions examined (56). This suggested that Fis has a regulatory role, however other factors are generally involved. An example of this is the demonstration that Fis bound to the *flh* regulatory region but no changes in expression were observed.

We also showed that Fis is a direct positive regulator of swarming motility through modulation of the lateral flagellum operon *laf* and the *scrABC* operon. The *scrABC* operon in *V. parahaemolyticus* controls the transition between swarming motility and adhesion to a surface by altering gene expression of *laf* and *cps* operons (46, 58). Together, ScrA, ScrB, and ScrC modulate the level of c-di-GMP in the cell, a secondary messenger that controls numerous downstream processes. More specifically, ScrC contains both GGDEF and EAL enzymatic activity, making it a unique bifunctional enzyme (58, 59). ScrA produce the extracellular S-signal which represses CPS gene expression. In the presence of ScrA, which interacts with ScrB, ScrC acts as a phosphodiesterase to degrade c-di-GMP. High levels of c-di-GMP promote CPS production, while low levels of c-di-GMP promote swarming motility (48, 59). In the Δ*fis* mutant, we observed down regulation of both the *laf* and *scrABC* operons in the GFP reporter assay, indicating that Fis is a positive regulator of these operons. We suggest that the Δ*fis* mutant swarming defect is through down-regulation of both the *laf* and *scrABC* operons and through the derepression of *opaR*, which is a repressor of swarming motility and an activator of CPS production (**Fig. 10**). Therefore, Fis integrates both the QS and surface sensing signals by positively regulating the *qrrs* and *scrABC* to induce swarming motility and repress CPS formation (**Fig. 10**).

**FIG 10.**
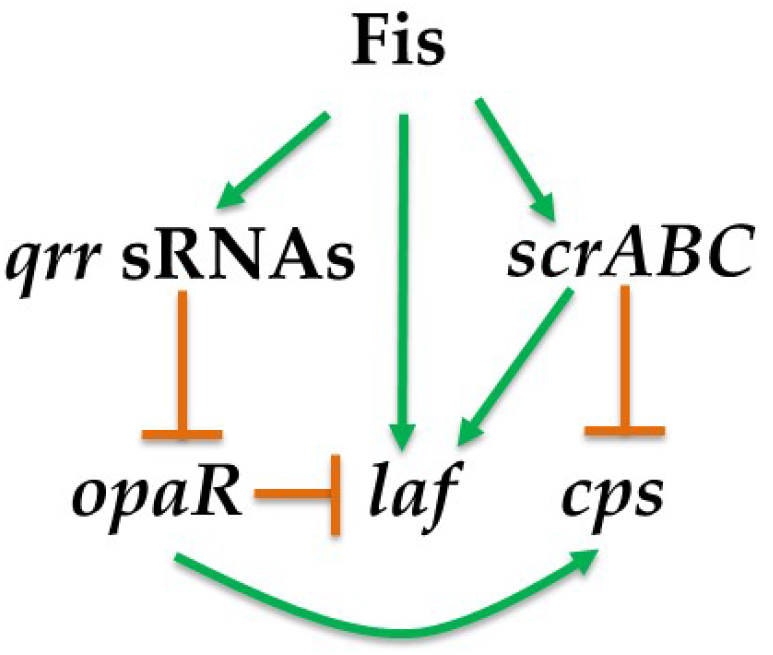
Model of Fis integration of QS and surface sensing pathways to control motility. Model shows control of swarming (*laf*) and CPS (*cps*) production in *V. parahaemolyticus*. Green arrows show direct positive regulation and gold hammers show direct negative regulation.

In *Salmonella enterica*, Fis was shown to negatively regulate genes contributing to virulence and metabolism in the mammalian gut (9). It was demonstrated that Fis was a negative regulator of acetate metabolism, biotin synthesis, fatty acids metabolism and propanediol utilization, amongst others, and speculated that this could be important for intestinal colonization and/or systemic infection (9). In *V. parahaemolyticus*, the *in vivo* competition assays in the streptomycin pretreated mouse model using Δ*fis* strain and WBWLacZ, showed that the Δ*fis* strain was outcompeted by wild type, demonstrating that Fis is required for *in vivo* fitness. In addition, *in vitro* growth competition assays in intestinal mucus and mucus components demonstrated that Δ*fis* was also outcompeted by wild type. We speculate that the defect in colonization of the mutant is due to its inability to efficiently utilize nutrient sources. Carbon metabolism was previously implicated as important for colonization of *V. parahaemolyticus* in streptomycin pretreated adult mouse model (41, 45). In summary, Fis modulates expression of genes involved in QS, motility, and metabolism in *V. parahaemolyticus* and it will be of interest to determine the different mechanisms used to modulate expression by this NAP.

## MATERIALS AND METHODS

### Bacterial strains, media, and culture conditions

All strains and plasmids used in this study are listed in **Table S1**. A streptomycin-resistant clinical isolate *V. parahaemolyticus* RIMD2210633 was used in this study. Unless stated otherwise, all *V. parahaemolyticus* strains were grown in lysogeny broth (LB) medium (Fischer Scientific, Pittsburgh, PA) containing 3% NaCl (LBS) at 37°C with aeration or M9 medium media (Sigma Aldrich, St. Louis, MO) supplemented with 3% NaCl (M9S). Antibiotics were added to growth media at the following concentrations: Ampicillin (Amp), 100 μg/ml, streptomycin (Sm), 200 μg/ml, tetracycline (Tet), 1 μg/mL, and chloramphenicol (Cm), 12.5 μg/ml when required.

### Construction of Δ*fis* mutant in *V. parahaemolyticus* RIMD2210633

Splicing by overlap extension (SOE) PCR and an allelic exchange method (60) were used to construct an in-frame, non-polar deletion mutant of *fis* (VP2885) in *V. parahaemolyticus* RIMD2210633. Briefly, primers were designed using *V. parahaemolyticus* RIMD2210633 genomic DNA as a template. All primers used in this study are listed in **Table S2**. SOE PCR was conducted to obtain an 18 bp-truncated version of VP2885 (297-bp). The Δ*fis* PCR fragments were cloned into the suicide vector pDS132 (61) and named pDSΔ*fis*. pDSΔ*fis* was then transformed into *E. coli* strain β2155 λ*pir* (62), and conjugated into *V. parahaemolyticus* RIMD2210633. Conjugation was conducted by cross streaking both strains onto LB plate containing 0.3 mM DAP. The colonies were verified for single crossover via PCR. The colonies that had undergone a single crossover were grown overnight in LBS with no antibiotic added and plated onto LBS containing 10% sucrose to select for double crossover deletion mutant. The gene deletion was confirmed by PCR and sequencing.

### Phenotype assays

To observe CPS, Heart Infusion media containing 1.5 % agar, 2.5mM CaCl_2_, and 0.25% Congo red dye was used and incubated at 30°C. Biofilm assays were conducted using crystal violet staining. Cultures were grown overnight in LBS and the overnight grown culture (1:40 dilution) was then inoculated in LBS in a 96 well plate, static at 37°C. After 24 hours, the wells are washed with PBS, stained with crystal violet for 30 min and then accessed for biofilm formation. The biofilms are then dissolved in DMSO and OD_595_ was measured. Swimming assays were conducted in LB 2% NaCl with 0.6% agar and swarming assays were conducted in heart infusion (HI) media with 2% NaCl and 1.5% agar. To study swimming behavior, a single colony of the bacterium was stabbed into the center of the plate, and plates were incubated at 37°C for 24 hrs. For the swarming assay, plates were spot incubated on the surface of the media and grown at 30°C for 48 hrs.

### Bioinformatics analysis to identify putative Fis binding sites

The regulatory region of gene clusters of interest of *V. parahaemolyticus* RIMD210633 were obtained using NCBI nucleotide database. Virtual footprint, an online database, was used to identify putative Fis binding sites using the *E. coli* Fis consensus binding sequence (63). The 229-bp, 416-bp, 371-bp, 385-bp, 153-bp, 385-bp, and 545 bp DNA regions upstream of *flhA* (VP2235-VP2231), *lafB (*VPA1550-VPA1557), *araB* (VPA1674), *nagB* (VPA0038), *gntK* (VP0064), and *scrABC* (VPA1513) respectively, were used as an input for Fis binding. The regulatory regions of *qrr1* (193-bp), *qrr2* (338-bp), *qrr3* (162-bp), *qrr4* (287-bp), and *qrr5* (177-bp) were also used as an input. Default settings were used to obtain putative Fis binding sites. A 130-bp sequence of VPA1424 regulatory region was used as a negative control and a 229-bp sequence of *gyrA* regulatory region was used a positive control.

### Fis protein purification

Fis was purified using a method previously described with modifications as necessary (45, 64). Briefly, Fis was cloned into the pMAL-c5x expression vector in which a 6X His tag maltose binding protein (MBP) was fused to *fis* separated by a tobacco etch virus (TEV) protease cleavage site (65). Primer pair FisFWDpMAL and FisREVpMAL (**Table S2**) and *V. parahaemolyticus* RIMD2210633 genomic DNA were used to amplify *fis* (VP2885). The *fis* PCR product along with purified pMAL-c5x, were digested with NcoI and BamHI ligated with T4 ligase and transformed into DH5α. The vector pMAL-c5x*fis* was purified, sequenced, and then transformed into *E. coli* BL21 (DE3). A 10 mL portion of *E. coli* BL21 pMAL-c5x*fis* overnight cultures were used to inoculate 1 L of fresh LB supplemented with 100 μg/ml ampicillin and 0.2% glucose and was grown at 37° C until the OD reached 0.4, at which point, the culture was induced by adding 0.5 mM IPTG. The cells were grown overnight at 18° C. Cells were pelleted at 5000 rpm and resuspended in 15 ml of column buffer (50 mM sodium phosphate, 200 mM NaCl, pH 7.5) supplemented with 0.5 mM benzamidine, and 1mM phenylmethylsulphonyl fluoride (PMSF). Bacterial cells were lysed using a microfluidizer, spun down at 15000 RPM for 60 min, and the supernatant was collected. The supernatant was passed through a 20 ml amylose resin (New England BioLabs) and washed with 10 column volumes (CVs) of column buffer. Fis fused with 6X His-MBP and was then eluted with three CVs column buffer supplemented with 20 mM maltose. Using 6XHis-TEV protease (1:10, TEV:protein in 50mM sodium phosphate, 200mM NaCl, 10mM imidazole, 5mM BME, pH 7.5) the fused protein was cleaved at TEV cleavage site. The cleaved fused protein was adjusted to 20 mM imidazole and run through immobilized metal affinity chromatography (IMAC) column using HisPur Ni-NTA resin to remove the cleaved 6XHis-MBP and the 6XHis-TEV protease. Mass spectrometry was performed to confirm Fis protein molecular weight and SDS-PAGE was conducted to determine its purity.

### Electrophoretic Mobility Shift Assays (EMSA)

Both a 138-bp and 152-bp fragment of P*araB* (VPA1674, arabinose catabolism), a 154-bp fragment of P*nagB* (VPA0038, glucosamine catabolism), 138-bp fragment of P*gntK* (VP0064, gluconate catabolism), a 161-bp fragment of P*flhA* (VP2235-VP2231), a 244-bp fragment of P*lafB* (VPA1550-VPA1557), and a 545-bp fragment of P*scrABC* (VPA1513-VPA1515) regulatory regions were used as probes in electrophoretic mobility shift assays (EMSA). Additionally, the full regulatory regions of all five *qrrs* were also analyzed for binding of Fis. A 193-bp fragment of P*qrr1*, a 338-bp fragment of P*qrr2*, a 162-bp fragment of P*qrr3,* a 287-bp fragment of P*qrr4,* and a 177-bp fragment of P*qrr5* regulatory regions were used as probes. A 130-bp probe of VPA1424 regulatory region was used as a negative control and a 229-bp probe of *gyrA* regulatory region as a positive control. The fragments were PCR amplified using Phusion Hifidelity Polymerase in 50 μl reaction mixture using respective primers sets listed in **Table S2** and *V. parahaemolyticus* RIMD2210633 genomic DNA as template. Various concentration of purified Fis were incubated with 30 ng of target DNA in binding buffer (10 mM Tris, 150 mM KCL, 0.1 mM dithiothreitol, 0.1 mM EDTA, 5% PEG, pH7.4) for 20 min at room temperature. A native acrylamide 6% gel was prepared and pre-run for 2 hours (200 V at 4°C) with 1x Tris-acetate-EDTA (TAE) buffer, and then 10 μl of the target DNA-protein mixture was loaded into consecutive lanes. The gel was run at 200V for 2 hours in 1X TAE buffer at 4°C, which was then stained in an ethidium bromide bath (0.5 μg/ml) for 20 min and imaged.

### Reporter assays

GFP reporter assays were conducted in *V. parahaemolyticus* RIMD2210633 and Δ*fis* strains. Reporter plasmids were constructed with the full regulatory regions of motility genes *flhAFG* and *lafBCDELSTI* and metabolic genes *araBDA*, *nagB* and *gntK* upstream of a promoterless *gfp* gene, as previously described (66). Briefly, primers were designed to amplify the regulatory region upstream of each gene or gene cluster with primer pairs listed in **Table S2**. Each amplified regulatory region was then ligated with the promoterless parent vector pRU1064 (67), which had been linearized prior with SpeI, using NEBuilder High Fidelity (HiFi) DNA Assembly Master Mix (New England Biolabs, Ipswich, MA) via Gibson Assembly Protocol (68). Overlapping regions for Gibson Assembly are indicated in lower case letters in the primer sequence in **Table S2**. Reporter plasmid P_*flhA-gfp*_ encompasses 269-bp of the regulatory region upstream of *flhA*. Reporter plasmid P_*lafB-gfp*_ encompasses 456-bp of the regulatory region upstream of *lafB*. Reporter plasmid P_*araB-gfp*_ encompasses 411-bp of the regulatory region upstream of *araB*. Reporter plasmid P_*nagB-gfp*_ encompasses 434-bp of the regulatory region upstream of *nagB*. Reporter plasmid P_*gntK-gfp*_ encompasses 193-bp of the regulatory region upstream of *gntK*. Reporter plasmid P_*scrABC-gfp*_ encompasses 545-bp of the regulatory region upstream of *scrABC.* Additionally, the full regulatory region of *qrr1* to *qrr5* were amplified and cloned into the pRU1064 reporter plasmid. The plasmids were transformed into *E. coli* Dh5α, purified and sequenced. Plasmids were then conjugated into wild type and the Δ*fis* mutant for further analysis.

Strains were grown overnight with aeration at 37°C in LBS with tetracycline (1 μg/ml). Cells were then pelleted, washed two times with 1X PBS, diluted 1:100 in LBS. Strains containing the P*qrr* and P*opaR* reporters were grown to low cell density (0.4-0.45 OD), washed two times with 1X PBS, and resuspended to a final OD of 1.0 before measuring relative fluorescence. The metabolism gene reporters were grown for 20 hrs with antibiotic selection. P_*araB-gfp*_ was grown in LBS supplemented with 10 mM D-arabinose, P_*nagB-gfp*_ was grown in LBS supplemented with 10 mM D-glucosamine, P_*gntK-gfp*_ was grown in LBS supplemented with 10 mM D-gluconate. Cells were pelleted and resuspended in 1X PBS. The pRUP*lafB* reporter assay was performed using cells grown on heart-infusion (HI) plates for 16 h. Colonies were scraped from the plate and resuspended in 1xPBS to a final OD_595_ around 0.5. GFP fluorescence was measured with excitation at 385 and emission at 509 nm in black, clear-bottom 96-well plates on a Spark microplate reader with Magellan software (Tecan Systems Inc.). Specific fluorescence was calculated for each sample by normalizing relative fluorescence to OD_595_. At least two biological replicates, in triplicate, were performed for each assay. Statistics were calculated using an unpaired Student’s t-test.

### *In vitro* growth competition assays

*In vitro* growth competition assays were performed by diluting an inoculum 1:50 into LBS broth, and separately in M9S supplemented with 10mM of individual carbon sources, D-glucose, L-arabinose, L-ribose, D-gluconate, D-glucosamine, and N-acetylglucosamine (NAG). In all cases, the culture was incubated at 37°C for 24 h, serially diluted and plated on LBS plus streptomycin and 5-bromo-4-chloro-3-indolyl-B-D-galactoside (X-gal). The competitive Index (CI) was determined using the following equation: CI=ratio out (Δ*fis* /WBWlacZ) / ratio in (Δ*fis*/WBWlacZ). A CI of < 1 indicates WBWlacZ outcompetes the Δ*fis* mutant, a CI of > 1 indicates that the Δ*fis* mutant outcompetes WBWlacZ. The ratio of Δ*fis* to WBWlacZ in the inoculum mixture is termed as “Ratio in” and the ratio of Δ*fis* to WBWlacZ colonies recovered from the mouse intestine is referred as “Ratio out.”

### *In vivo* colonization competition assays

All mice experiments were approved by the University of Delaware Institutional Animal Care and Use Committee. A β-galactosidase knock in *V. parahaemolyticus* RIMD2210633 strain, WBWlacZ, which was previously shown to grow similarly to wild type *in vitro* and *in vivo*, was used for all the competition assays (41, 50, 69). Inoculum for competition assays was prepared using overnight cultures of WBWlacZ and Δ*fis* diluted into fresh LBS media and grown for four hours. These exponential phase cultures were then pelleted by centrifugation at 4,000 × g, washed and resuspended in PBS. One mL of WBWlacZ and one mL of Δ*fis* were prepared, corresponding to 1 x 10^10^ CFU of each strain, based on the previously determined OD and CFU ratio. A 500 μl aliquot of Δ*fis* was combined with 500 μl of the WBWlacZ, yielding a total bacterial concentration of 1 x 10^10^ CFU/mL. The inoculum was serially diluted and plated on LBS agar plate supplemented with 200 μg/ml streptomycin and 8 μg /ml of X-gal to determine the exact ratio of the inoculum. Male C57BL/6 mice, aged 6 to 10 weeks were housed under specific-pathogen-free conditions in standard cages in groups (5 per group) and provided standard mouse feed and water *ad libitum*. Pretreatment of mice with streptomycin was performed as previously described (50, 69). Mice were inoculated with 100 μl of the bacterial suspension and 24 hours post-infection, mice were sacrificed, and the entire gastrointestinal tract was harvested. Samples were placed in 8 mL of sterile 1x PBS, mechanically homogenized and serially diluted in 1x PBS. Diluted samples were plated for CFU’s on LBS, supplemented with streptomycin and X-gal for a blue (WBWlacZ) versus white (Δ*fis*) screen of colonies after incubation at 37°C overnight. The competitive index (CI) was determined as described above.

## ACKNOWLEDGEMENTS

This research was supported in part by a National Science Foundation grant (award IOS-1656688) to E.F.B. J.G.T was funded by a University of Delaware graduate fellowship award. We thank members of the Boyd Group for constructive feedback on the manuscript.

